# Epigenetic programming of host lipid metabolism associates with resistance to TST/IGRA conversion after exposure to *Mycobacterium tuberculosis*

**DOI:** 10.1101/2024.02.27.582348

**Authors:** Kimberly A Dill-McFarland, Jason D Simmons, Glenna J Peterson, Felicia K Nguyen, Monica Campo, Penelope Benchek, Catherine M Stein, Tomas Vaisar, Harriet Mayanja-Kizza, W Henry Boom, Thomas R Hawn

## Abstract

*Mycobacterium tuberculosis* (Mtb) exposure leads to a range of outcomes including clearance, latent TB infection (LTBI), and pulmonary tuberculosis (TB). Some heavily exposed individuals resist tuberculin skin test (TST) and interferon gamma release assay (IGRA) conversion (RSTR), which suggests that they employ IFNγ-independent mechanisms of Mtb control. Here, we compare monocyte epigenetic profiles of RSTR and LTBI from a Ugandan household contact cohort. Chromatin accessibility did not differ between uninfected RSTR and LTBI monocytes. In contrast, methylation significantly differed at 174 CpG sites and across 63 genomic regions. Consistent with previous transcriptional findings in this cohort, differential methylation was enriched in lipid and cholesterol associated pathways including in the genes APOC3, KCNQ1, and PLA2G3. In addition, methylation was enriched in Hippo signaling, which is associated with cholesterol homeostasis and includes CIT and SHANK2. Lipid export and Hippo signaling pathways were also associated with gene expression in response to Mtb in RSTR as well as IFN stimulation in monocyte-derived macrophages (MDMs) from an independent healthy donor cohort. Moreover, serum-derived HDL from RSTR had elevated ABCA1-mediated cholesterol efflux capacity (CEC) compared to LTBI. Our findings suggest that resistance to TST/IGRA conversion is linked to regulation of lipid accumulation in monocytes, which could facilitate early Mtb clearance among RSTR subjects through IFNγ-independent mechanisms.

**IMPORTANCE:** Tuberculosis (TB) remains an enduring global health challenge with millions of deaths and new cases each year. Despite recent advances in TB treatment, we lack an effective vaccine or a durable cure. While heavy exposure to *Mycobacterium tuberculosis* often results in latent TB latent infection (LTBI), subpopulations exist who are either resistant to infection or contain Mtb with IFNγ-independent mechanisms not indicative of LTBI. These resisters provide an opportunity to investigate mechanisms of TB disease and discover novel therapeutic targets. Here, we compare monocyte epigenetic profiles of RSTR and LTBI from a Ugandan household contact cohort. We identify methylation signatures in host lipid and cholesterol pathways with potential relevance to early TB clearance before the sustained IFN responses indicative of LTBI. This adds to a growing body of literature linking TB disease outcomes to host lipids.

## INTRODUCTION

Tuberculosis (TB) remains one of the leading causes of single-agent infectious disease death worldwide with over 1 in 1000 people developing new TB and 1.4 million deaths annually (1). Individuals infected with *Mycobacterium tuberculosis* (Mtb), the causative agent of TB, exhibit a range of clinical phenotypes from latent Mtb infection (LTBI) to pulmonary TB disease. LTBI, defined as a positive tuberculin skin test (TST) or interferon (IFN)-γ release assay (IGRA) with no clinical or radiographic evidence of TB disease, describes as much as a quarter of the world’s population (2). Some individuals resist TST/IGRA conversion (RSTR) despite prolonged and high-level Mtb exposure (3–6). Many factors likely contribute to resistance to TST/IGRA conversion including genetics, previous *Mycobacterium* infections, lung function, and others (7). We and others previously found differences in immunologic profiles of RSTR and LTBI subjects (5), including transcriptional signatures in monocytes (8, 9). Further investigation of RSTR populations may yield novel insights into host responses to Mtb and identify IFNγ-independent correlates of protection or targets for TB therapeutics.

Epigenetic mechanisms may regulate transcriptional responses and contribute to RSTR mechanisms. Methylation has been associated with the risk of LTBI conversion (10, 11), anti-mycobacterial activity (12–14), and response to Bacille Calmette-Guérin (BCG), a vaccine for TB disease in children (15). As one of Mtb’s first targets in the lung, macrophages are poised to accomplish early Mtb clearance through cell intrinsic microbicidal pathways or by priming IFNγ-independent cellular responses, both of which may operative in the absence of TST/IGRA conversion (16, 17). Some macrophage responses to Mtb infection *in vitro* are modulated through methylation (18), and monocyte responses to BCG have been associated with chromatin remodeling (19). We also previously found that histone deacetylases (HDAC) were differentially expressed in RSTR monocytes compared to LTBI in a Ugandan household contact study (20) and that an HDAC3 inhibitor modulated macrophage signaling during Mtb infection (21). Together, these studies suggest that epigenetic “trained immunity” may regulate Mtb clearance without TST/IGRA conversion.

In this study, we further characterize immune pathways in a Ugandan RSTR cohort consisting of household contacts of pulmonary TB cases who remain TST/IGRA-negative over 8 - 10 years (3, 22). Genome-wide methylation and chromatin accessibility were assessed in unstimulated RSTR and LTBI monocytes. Orthogonal to RSTR transcriptional signatures identified previously (8, 9), we identify methylation signatures in lipid and cholesterol pathways with potential relevance to IFN responses and macrophage function during Mtb infection.

## RESULTS

### RSTR methylation differs in lipid and cholesterol associated genes compared to LTBI

We examined chromatin accessibility and methylation in monocytes from RSTR and LTBI individuals (Table 1). RSTR and LTBI groups did not differ by sex, age, body mass index (BMI), BCG vaccination scar, TB exposure score a enrollment (3), or relatedness as measured by total first- and third-degree or closer relationships between each individual and all other individuals in the dataset (P > 0.05). All participants were HIV negative.

**Table 1.**
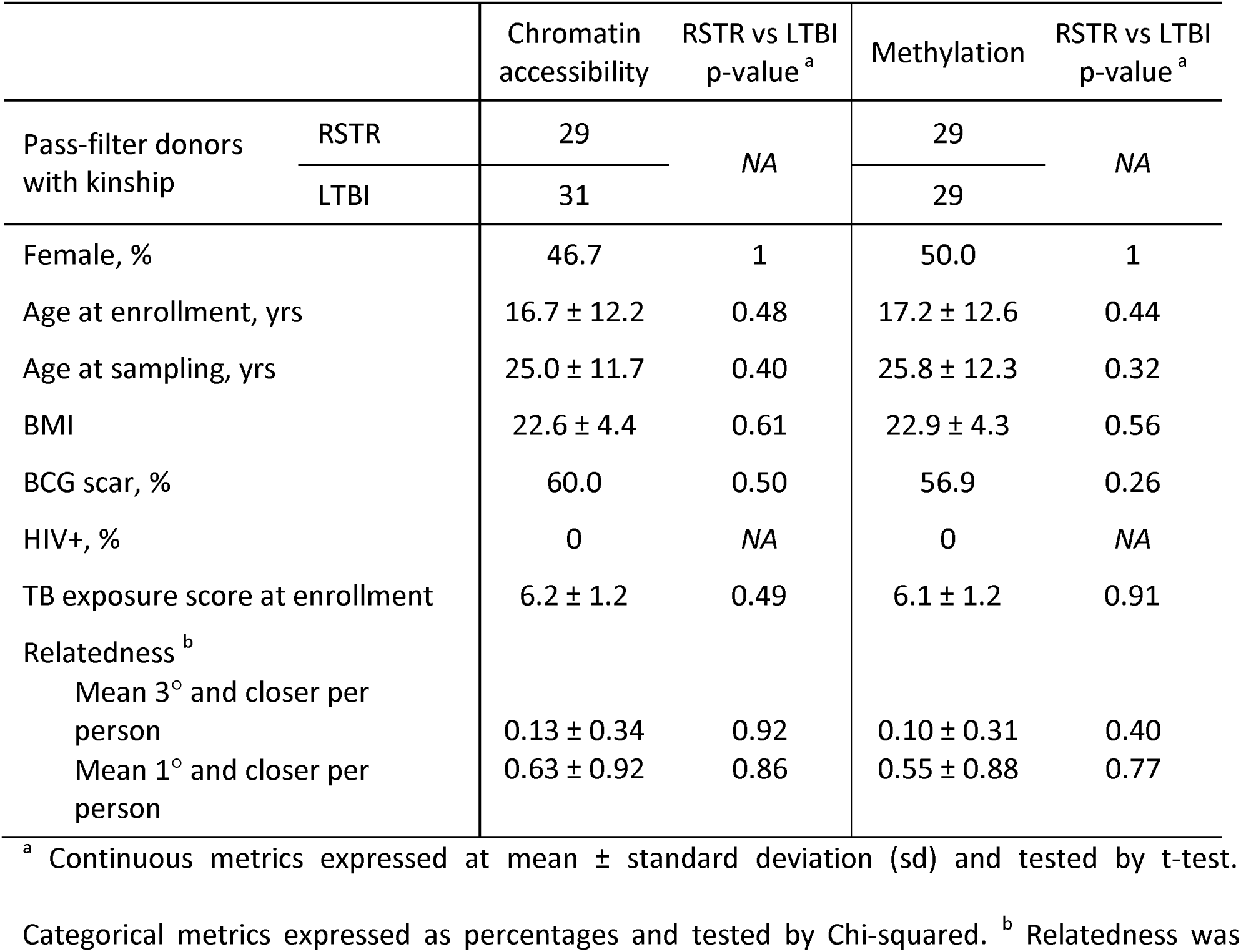
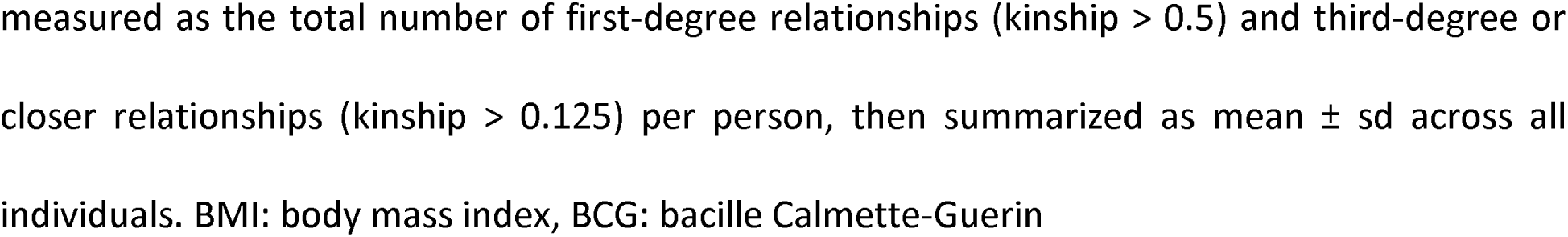

|  |  | Chromatin accessibility | RSTR vs LTBI p-value <sup>a</sup> | Methylation | RSTR vs LTBI p-value <sup>a</sup> |
| --- | --- | --- | --- | --- | --- |
| Pass-filter donors with kinship | RSTR | 29 | NA | 29 | NA |
|  | LTBI | 31 |  | 29 |  |
| Female, % |  | 46.7 | 1 | 50.0 | 1 |
| Age at enrollment, yrs |  | 16.7 ± 12.2 | 0.48 | 17.2 ± 12.6 | 0.44 |
| Age at sampling, yrs |  | 25.0 ± 11.7 | 0.40 | 25.8 ± 12.3 | 0.32 |
| BMI |  | 22.6 ± 4.4 | 0.61 | 22.9 ± 4.3 | 0.56 |
| BCG scar, % |  | 60.0 | 0.50 | 56.9 | 0.26 |
| HIV+, % |  | 0 | NA | 0 | NA |
| TB exposure score at enrollment |  | 6.2 ± 1.2 | 0.49 | 6.1 ± 1.2 | 0.91 |
| Relatedness <sup>b</sup> |  |  |  |  |  |
| Mean 3° and closer per person |  | 0.13 ± 0.34 | 0.92 | 0.10 ± 0.31 | 0.40 |
| Mean 1° and closer per person |  | 0.63 ± 0.92 | 0.86 | 0.55 ± 0.88 | 0.77 |

<sup>a</sup> Continuous metrics expressed at mean ± standard deviation (sd) and tested by t-test.
Categorical metrics expressed as percentages and tested by Chi-squared. <sup>b</sup> Relatedness was
measured as the total number of first-degree relationships (kinship > 0.5) and third-degree or closer relationships (kinship > 0.125) per person, then summarized as mean ± sd across all individuals. BMI: body mass index, BCG: bacille Calmette-Guerin

Epigenetic signals were tested using a linear mixed effects model for RSTR vs LTBI corrected for age, sex, and genetic kinship (Figure S1). Two differentially accessible regions (DAR) and 174 differentially methylated probes (DMP) were identified between RSTR and LTBI at FDR < 0.2 (Figure 1 A,B, Table S1). DMP were then used to define 63 differentially methylated regions (DMR) (FDR < 1E-70) (Figure 1C) each containing two to seven probes (Table S1). In total, 44 of the 63 DMR contained at least one DMP site (Table S1). Significant DAR annotated to one gene (CAMK1D) and an intergenic region. DMP annotated to 125 genes and DMR to 44 genes. Significant epigenetic sites included 40 DMPs and 18 DMRs annotated to promoter regions within 1500 bp of a transcription start site (TSS1500) (Figure S2). Together, these data suggest that RSTR and LTBI monocytes contain distinct epigenetic profiles long after their initial Mtb exposure.

**Figure 1.**
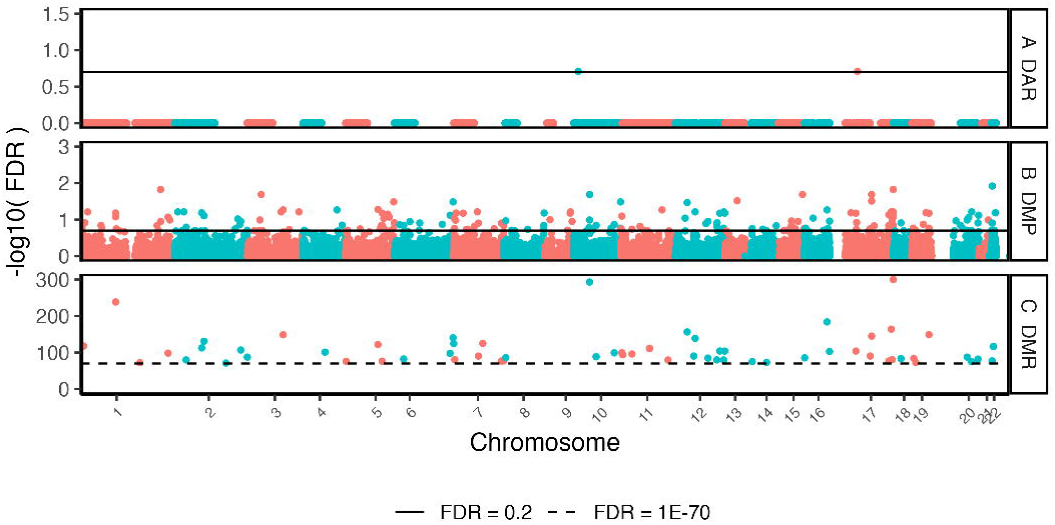
Monocyte epigenetic sites associated with RSTR vs LTBI. Genome-wide methylation and chromatin accessibility were assessed in unstimulated monocytes. RSTR and LTBI were compared using a mixed effects model corrected for age, sex, and kinship. (A) Differentially accessible chromatin regions (DAR, N = 40,919) were measured using log2 counts per million (CPM) reads in open chromatin peaks. (B) Differentially methylated probes (DMP, N = 728,380) and (C) differentially methylated DMRcate regions (DMR, N = 63) were measured using log2 M values. Significant hits are defined as FDR < 0.2 for DAR and DMP (solid line) or FDR < 1E-70 for DMR (dashed line). This resulted in 2 DAR, 174 DMP, and 63 DMR significantly different between RSTR and LTBI.

To assess for connections among the methylation genes, we used hypergeometric mean pathway enrichment analysis with Broad MSigDB gene-sets (23). DMR genes were enriched in 11 gene ontology (GO) pathways (FDR < 0.2 and k/K > 0.04, Table S2). No Hallmark gene-sets were significantly enriched, and neither database was enriched within DMP genes. The most highly enriched GO gene-sets for DMR were associated with high-density lipoprotein (HDL) remodeling, lipid export, fatty acid biosynthesis, and Hippo signaling (Figure 2A). Nine DMRs annotated to 8 unique genes (APOC3, CIT, KCNQ1, MCM2, PER3, PLA2G3, RPTOR, SHANK2) drove significant enrichments (Figure 2B, Table S3).

**Figure 2.**
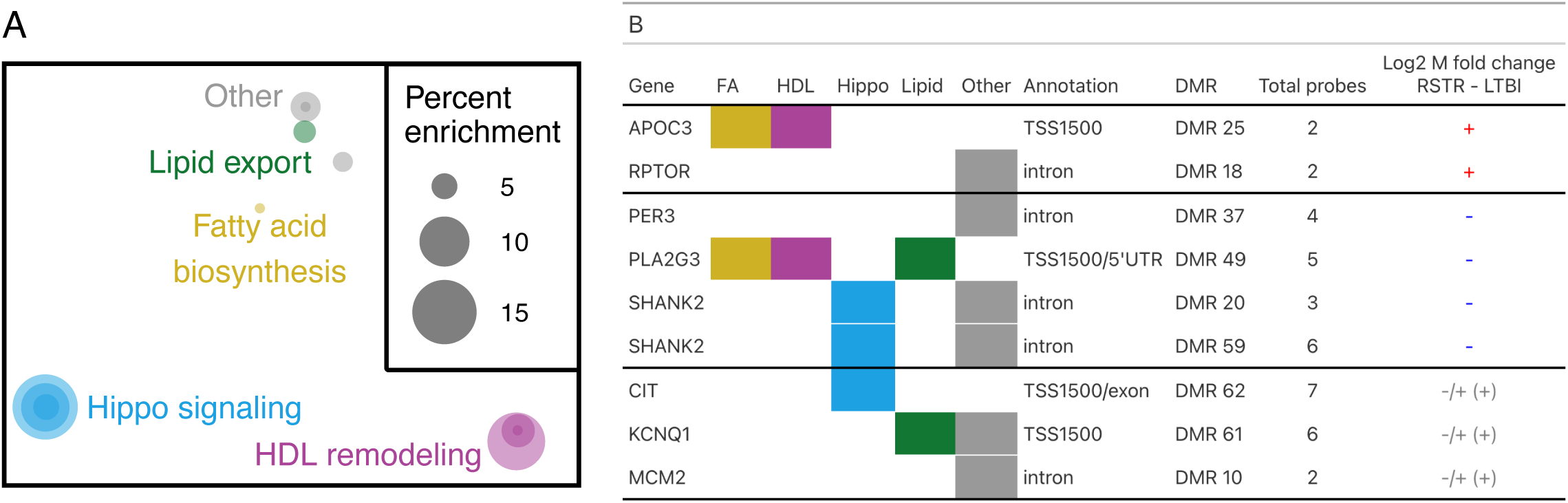
Pathways and genes with differentially methylated regions in RSTR and LTBI monocytes. Differentially methylated regions (DMR) were analyzed with hypergeometric mean pathway enrichment analysis with Broad MSigDB gene sets. (A) Gene ontology gene sets significantly enriched in DMR (FDR < 0.2 and k/K > 0.04). Gene sets visualized by semantic similarity in MDS space. Each dot is a gene set with size indicating percent enrichment (k/K * 100) and colors grouping similar terms. (B) DMR genes within significant GO gene sets. Colored boxes indicate presence in gene sets as in (A). Fold changes of probes in differentially methylated regions (DMR) are indicated by + positive for all probes, - negative for all probes, and -/+ variable across probes with the mean region trend in parentheses.

Among genes enriched in fatty acid, lipid, and HDL pathways (Figure 3A,B), APOC3, KCNQ1, and PLA2G3 each contained one DMR in their TSS1500 region. APOC3 had consistently higher methylation while PLA2G3 had consistently lower methylation as well as a significant DMP site (asterisk) in RSTR compared to LTBI (Figure 3A). In contrast, KCNQ1 had discordant DMR methylation and was driven by a single DMP site with higher methylation in RSTR (asterisk, Figure 3A). Among Hippo signaling associated genes (Figure 3C,D), CIT contained one TSS1500 DMR driven by a single DMP site with higher methylation in RSTR compared to LTBI (asterisk, Figure 3C). SHANK2 contained two intronic DMRs both with consistently lower methylation in RSTR compared to LTBI (Figure 3C,D). Together, these data suggest that methylation signatures differentiate RSTR and LTBI monocytes with profiles related to lipid and cholesterol processes.

**Figure 3.**
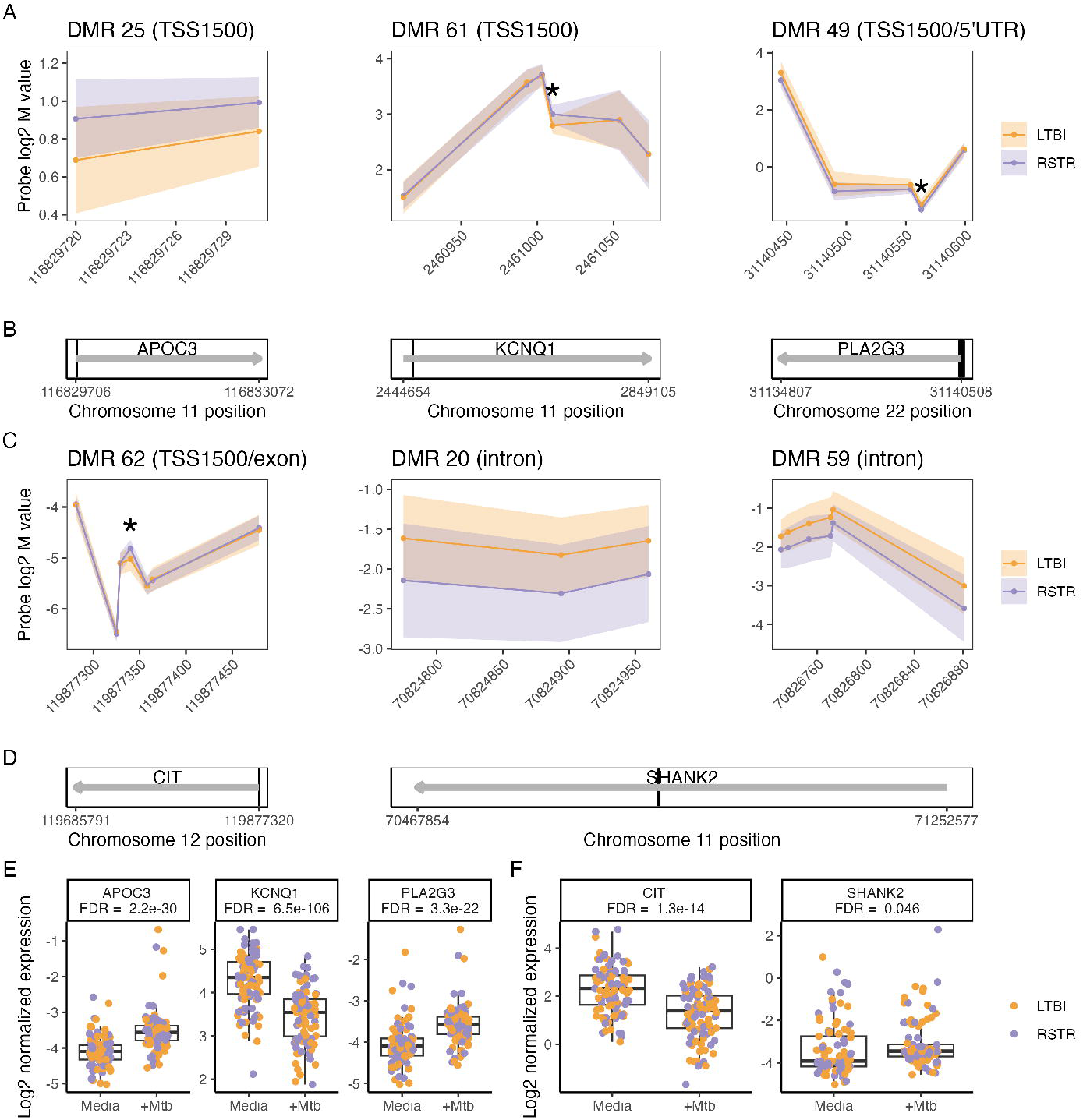
Methylation regions associated with significantly enriched gene sets in RSTR and LTBI monocytes. Genes annotated to differentially methylated regions (DMR, FDR < 1E-70) were assessed for enrichment in gene ontology (GO) gene sets. DMR associated with significantly enriched gene sets (FDR < 0.2) are grouped by (A,B,E) Ss or (C,D,F) Hippo signaling associated genes. The 3 DMR in other enriched pathways (Figure 2B) are not shown. (A,C) Log2 M values for each probe in a DMR are shown with standard deviation error shading for LTBI (orange) and RSTR (purple). Significant individual probes (DMP) are indicated with * (FDR < 0.2). (B,D) DMRs annotated to their nearest gene. Grey arrows indicate full gene transcripts, and black boxes are DMRs. SHANK2 includes DMR 20 and DMR 59 that are too close to be resolved in (D). (E,F) Log2 normalized gene expression in RSTR and LTBI monocytes with and without Mtb infection. FDR indicates the Mtb vs media comparison (FDR < 0.2). No genes were significant for the main effect RSTR vs LTBI or the interaction of Mtb:RSTR (FDR > 0.2).

Utilizing a previously published transcriptomics study of this same cohort (9), we next investigated the impacts of Mtb infection on gene expression related to methylation signatures. Among lipid and HDL-associated genes identified in the methylation analysis (Figure 2), APOC3 and PLA2G3 were significantly upregulated while KCNQ1 was downregulated in response to Mtb infection in monocytes (Figure 3E). For Hippo signaling genes, CIT was upregulated and SHANK2 was downregulated after Mtb infection (FDR < 0.05) (Figure 3F). None of these selected genes had differential expression between RSTR and LTBI groups (FDR > 0.2) (Figure 3E,F). Gene-set enrichment analysis (GSEA) further revealed that the lipid export gene-set was upregulated by Mtb infection in both RSTR and LTBI monocytes with the highest expression among Mtb-infected RSTR monocytes (P < 0.05, Figure 4A, Table S4). While GSEA suggested Hippo signaling was not significantly impacted by Mtb infection (FDR > 0.05), RSTR Mtb-infected monocytes had higher Hippo signaling gene expression compared to LTBI (FDR < 0.05, Figure 4A, Table S4). Thus, methylation signatures in uninfected monocytes correlate with Mtb-dependent gene expression patterns that distinguish RSTR from LTBI.

**Figure 4.**
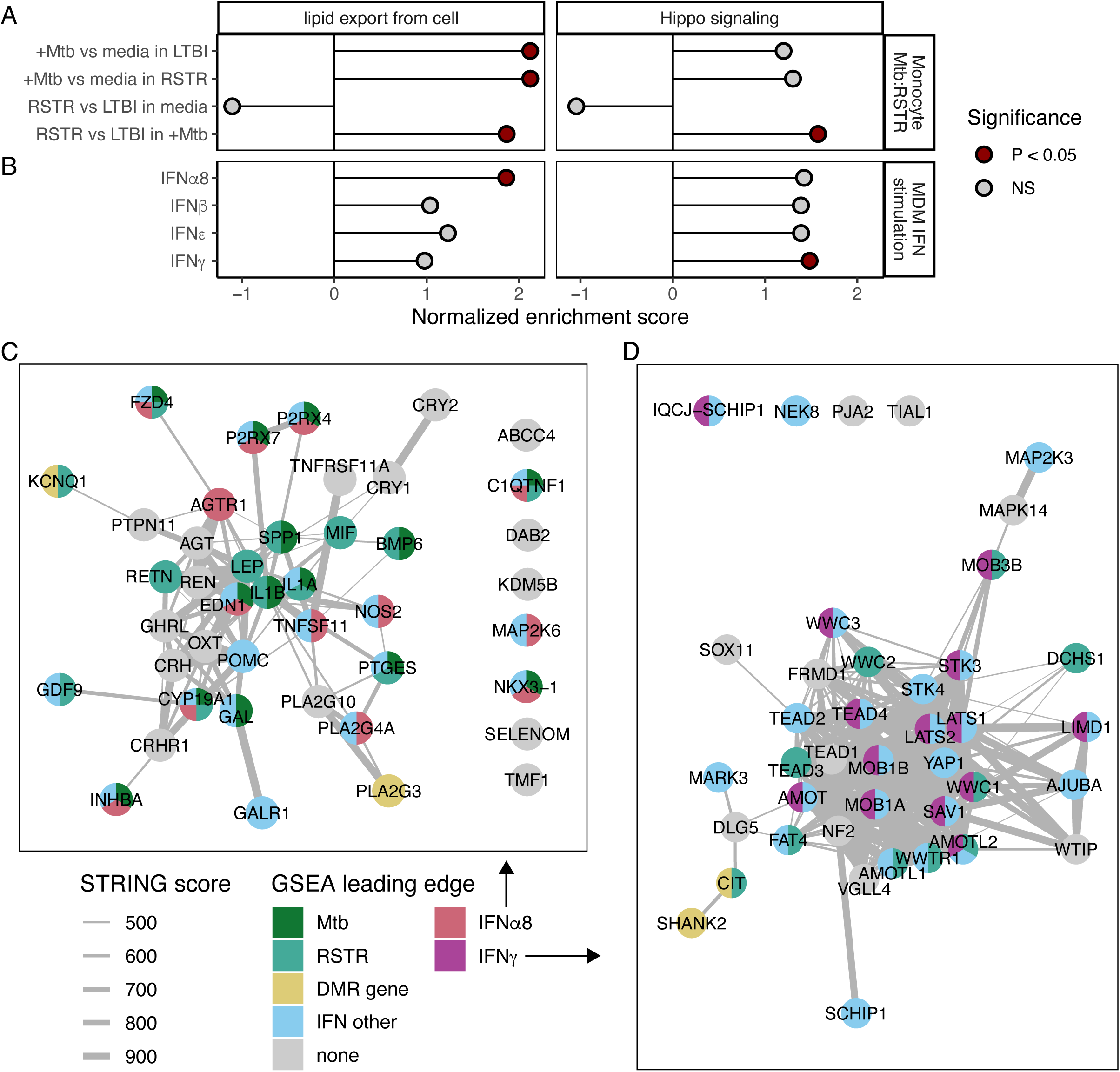
Mtb and IFN stimulated gene expression in gene sets enriched for RSTR methylation signature. Gene set enrichment analysis (GSEA) was performed for pathways that were significantly enriched for DMR annotated genes (N = 63 genes, Figure 2). Fold change estimates were used from mixed effects models run on one of two RNAseq data sets. (A) GSEA for Mtb and RSTR contrasts. Monocytes from RSTR and LTBI were infected with Mtb for 6 hours, and gene expression was modeled for the interaction of Mtb and RSTR status (Mtb:RSTR) corrected for age, sex, kinship, and sequencing batch. Contrast fold changes were compared for Mtb infection vs media within RSTR or LTBI as well as RSTR vs LTBI within media or Mtb infected groups. Significant enrichment is indicated in red (P < 0.05). (B) GSEA for IFN stimulation. MDMs from healthy donors were stimulated with type I and II interferons for 6 hrs. Gene expression was modeled for IFN vs media. (C) All genes in the GO gene set “lipid export from cell” are visualized in the STRING protein-protein interaction network. Color indicates leading-edge GSEA and significant DMR genes. Leading-edge genes include +Mtb vs media in RSTR and/or LTBI (Mtb), RSTR vs LTBI in Mtb-infected samples (RSTR), and IFNα8 stimulation vs media (IFNα8). Genes that were leading edge in non-significant IFNβ, IFNε, and IFNγ GSEA are grouped in “IFN other”. Genes annotated to a significant DMR are additionally colored (DMR gene). Edge width indicates STRING combined score with only scores > 400 shown. (D) GO “Hippo signaling” gene set genes are similarly represented, except leading-edge genes for IFNγ are colored and non-significant “IFN other” includes IFNα8, IFNβ, and IFNε.

### Interferon-specific lipid transcriptional responses in MDM

Given that RSTR and LTBI phenotypes are defined by Mtb-specific IFNγ responses, we next explored IFN-dependent gene expression in pathways associated with methylation. We looked for overlapping enrichment between methylation associated gene-sets described above and the transcriptional responses of an independent healthy cohort’s MDMs to Type I and Type II interferons (24). Among the 11 methylation-enriched GO gene-sets, lipid export was significantly up-regulated by IFNα8 stimulation in healthy MDMs (P < 0.05, Figure 4B, Table S4). We then utilized the STRING protein-protein interaction network to visualize all genes in the GO lipid export gene-set and determine overlap between methylation and transcriptional signatures (Figure 4C). The leading-edge genes for each enrichment analysis were colored to identify genes with differential methylation in RSTR (DMR gene) that likely drive expression changes in RSTR compared to LTBI (RSTR), in response to Mtb infection (Mtb), and/or in response to IFN stimulation (IFNα8, IFNγ, IFN other). As seen in the STRING network (Figure 4C), IFNα8 enrichment was driven by 13 leading-edge genes that mostly overlapped with leading-edge genes in the non-significant interferon stimulations (12/13, Figure 4D). This MDM IFNα8 leading-edge also overlapped with leading-edge genes for lipid export in response to Mtb (8/13) and RSTR compared to LTBI in Mtb infected monocytes (3/13). Together, these data suggest that epigenetic programming may impact lipid export pathways that lead to differences in RSTR and LTBI responses to Mtb infection and may be modulated by IFN induced responses.

In addition, the methylation-enriched Hippo signaling gene-set, which contributes to cholesterol homeostasis (25), was up-regulated by IFNγ (P < 0.05, Figure 4B). As visualized in the STRING network (Figure 4D), Hippo signaling in MDMs had 14 leading-edge genes that mostly overlapped with nonsignificant IFN stimulations (12/14) and somewhat overlapped with the RSTR compared to LTBI leading-edge in Mtb infected monocytes (3/13) (Figure 4D). Overall, these data indicate that epigenetically programmed pathways related to Hippo and cholesterol may be impacted by IFNγ induced responses that define RSTR and LTBI phenotypes.

### Cholesterol efflux capacity of HDL differs between RSTR and LTBI

To examine lipid-dependent mechanisms of RSTR clearance of Mtb, we next measured serum HDL particle concentrations and sizes as well as cholesterol efflux capacity (CEC) of serum HDL from RSTR and LTBI individuals. Neither HDL particle concentrations nor sizes (xsHDL to xlHDL) differed between unstimulated RSTR and LTBI serum (Figure S3, Table S5, FDR > 0.2). RSTR and LTBI serum HDL had similar total CEC with cAMP activation to stimulate efflux in mouse J774 macrophages (6.1 ± 1.9 %, P > 0.05, Figure 5A, Table S5). In contrast, ABCA1-specific CEC in BHK cells was higher with RSTR serum HDL compared to LTBI (Δ LTBI = 6.2 ± 2.1 %, Δ RSTR = 7.5 ± 2.8 %, P = 0.02) with no differences in unstimulated cells (Figure 5B, Table S5). Moreover, investigation of the previously published transcriptomics study of this same cohort (9) revealed differential expression of ABCA1 in monocytes upon Mtb infection with RSTR displaying significantly higher induction compared to LTBI (Figure 5C). Together, the complimentary effects of increased CEC and ABCA1 may reduce intracellular cholesterol levels during infection and contribute to IFNγ-independent Mtb control in RSTR.

**Figure 5.**
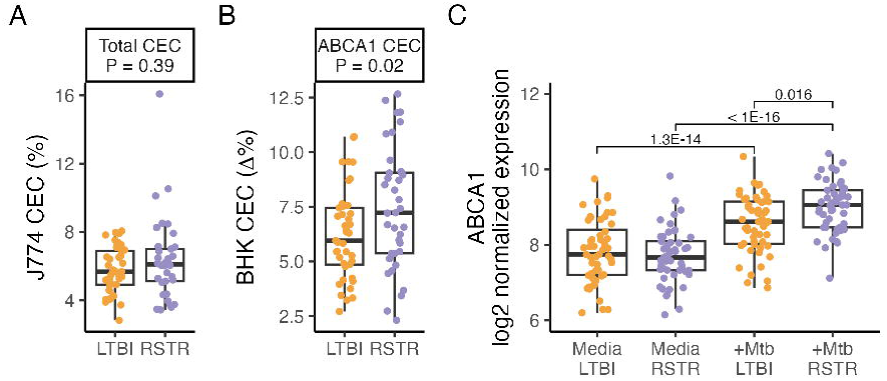
Total and ABCA1-specific cholesterol efflux capacity in RSTR and LTBI. Cholesterol efflux capacity (CEC) was measured in serum-derived HDL from RSTR and LTBI participants. Serum HDL was incubated for 4 hours with unstimulated or stimulated [^3^H]-cholesterol labeled cells. CEC was calculated as percent cholesterol in media versus total (media + cell pellet). RSTR and LTBI were compared using a linear model corrected for age sex, and body mass index (BMI). (A) Total CEC from J774 cells was measured after exposure to RSTR and LTBI serum HDL. Macrophages were stimulated with cAMP to increase CEC. (B) CEC in ABCA1-transfected BHK cells with RSTR and LTBI HDL. ABCA1-transfected BHK cells were stimulated with media or mifepristone to induce ABCA1 expression. ABCA1-specific CEC was calculated as stimulated (ABCA1 expressing) BHK minus unstimulated. (C) ABCA1 gene expression in monocytes with and without Mtb infection ^9^. RSTR and LTBI monocytes were cultured with and without Mtb infection for 6 hrs. Gene expression was modeled for the interaction between Mtb infection and RSTR status (Mtb:RSTR). Significant Mtb:RSTR contrasts are labeled by FDR (FDR < 0.2).

## DISCUSSION

We assessed monocyte methylation and chromatin accessibility signatures in latent tuberculosis infection (LTBI) and IGRA/TST resistant individuals (RSTR). We found differential methylation in genes related to high-density lipoprotein (HDL) remodeling, lipid export, and fatty acid metabolism. In addition, ABCA1-mediated HDL cholesterol efflux capacity (CEC) was higher in RSTR compared to LTBI. These results contribute to the growing body of literature supporting the importance of host lipid and fatty acid metabolism in Mtb infection and disease progression (8, 26).

RSTR had distinct methylation patterns compared to LTBI in genes associated with fatty acid metabolism and lipid transport. These genes included the phospholipase PLA2G3 (27) and lipase repressor APOC3 (28), which associate with HDL and impact intracellular lipid availability. Specifically, an overabundance of APOC3 may reduce HDL-mediated CEC, potentially through competitive binding to the scavenger receptor SCARB1 (29). In addition, the lipid-mediated voltage-gated potassium channel KCNQ1 contained methylation signatures in RSTR. Several KCNQ1 mutations are associated with plasma lipid accumulation (30), and KCNQ1’s antisense lncRNA, KCNQ1OT1, is associated with reduced CEC and increased lipid accumulation in macrophages specifically (31). These lipase and lipid transport activities impact free fatty acid (FFA) availability and contribute to the accumulation of lipids. This can lead to lipid-laden, or foamy, macrophages which are a permissive environment for Mtb persistence (32–34). In fact, PLA2G3 directly promotes foamy macrophage accumulation in mice (27). The importance of FFAs in the RSTR phenotype is further supported by previous work in this Ugandan as well as an independent South African cohort, where gene-sets associated with FFA metabolism were more highly expressed in unstimulated RSTR monocytes compared to LTBI (35). Finally, lipid transport after Mtb infection may be further regulated by IFN signaling, a defining feature of RSTR, as indicated by upregulation of lipid export gene expression in response to Mtb infection in RSTR monocytes and to IFNα8 in monocyte-derived macrophages (MDMs). Thus, differences in RSTR methylation in FFA metabolism and lipid transport genes correlate with transcriptional signatures and suggest that epigenetic programming may control lipid accumulation, thus ultimately preventing Mtb persistence.

The RSTR phenotype may be additionally mediated by cholesterol as indicated by the previously discussed lipid-associated genes (PLA2G3, APOC3, KCNQ1) as well as additional methylation hits related to Hippo signaling (CIT, SHANK2). Cholesterol represents an important fatty acid for Mtb as it is a preferred carbon source (32, 34) and contributes to macrophage responses to Mtb including membrane lipid rafts permissive to Mtb uptake (36), modulation of cellular function, and development of foamy macrophages (32, 33, 37, 38). Hippo signaling is involved in pleiotropic activities related to cell proliferation, differentiation, and survival (25) including in macrophages (39). Among these functions, Hippo signaling regulates cholesterol levels through sterol regulatory element-binding proteins (SREBPs) (25). We previously identified enrichments in IFNγ signaling that discriminated LTBI from RSTR monocytes (8, 9), an expected finding based on categoric IGRA responses (3, 22). Paradoxically, Hippo signaling expression was increased with IFNγ stimulation in healthy MDMs but associated with Mtb-infected RSTR monocytes. This suggests that Hippo signaling is activated among RSTR and may provide protection against Mtb infection in the absence of IFNγ (3, 22). We further investigated RSTR links to cellular cholesterol by measuring CEC. RSTR serum HDL resulted in higher ABCA1-mediated cholesterol efflux compared to LTBI, thus supporting that RSTR are better able to control intracellular cholesterol accumulation. Together, these findings support a hypothesis that Mtb growth is restricted among RSTR monocytes due to enhanced CEC that leads to decreased intracellular (40) or membrane (36) cholesterol with CEC potentially regulated by ABCA1 and/or Mtb-induced Hippo signaling as a result of specific methylation signatures.

Our study has several limitations. The peripheral blood monocytes investigated in this study represent an important cell population relevant to TB disease. However, epigenetic profiles in lung resident cell populations such as alveolar macrophages may reveal additional markers of the RSTR phenotype. Additionally, while methylation is known to alter gene expression, the impacts of hyper- vs hypomethylation are not clear for gene expression directionality or association with disease (41, 42). Therefore, a more complete analysis of RSTR with paired methylation and gene expression in Mtb-infected cells is needed to elucidate the impacts of methylation on Mtb-relevant gene expression. This design may also reveal differences in chromatin accessibility not captured in the current dataset from uninfected cells. While methylation remains relatively stable over time, changes can occur on the time scale of years to decades (43). This RSTR cohort was defined by long-term follow-up spanning up to 10 years. Thus, the epigenetic markers of this stringently defined RSTR phenotype that was sampled long after documented exposure may not fully capture mechanisms relevant to Mtb clearance at the time of exposure. Finally, these findings are based on a single RSTR cohort, and epigenetic inquiry into additional cohorts with varied RSTR definitions is needed to confirm findings applicable to a larger population.

Overall, these results indicate that RSTR have distinct epigenetic programming related to lipid and cholesterol transport in monocytes. Differentially methylated genes in RSTR and LTBI likely lead to differences in lipid accumulation in monocytes and this, in turn, may alter Mtb outcomes and contribute to IFNγ independent control of Mtb in RSTR. Together with previous work in this cohort (8, 9, 44), this study supports the role of lipid and fatty acid metabolism in defining the Ugandan RSTR cohort as well as provides novel targets for the development of TB treatment.

## METHODS

### Cohort

Individuals with culture-positive pulmonary tuberculosis (TB) were recruited as part of the Kawempe Community Health Study in Kampala, Uganda from 2002 to 2012 as previously described (3, 22). Household contacts of index cases were initially followed for 2 years with serial tuberculin skin test (TST) monitoring (22). A subset of individuals were retraced from 2014 and 2017 and re-assessed by TST as well as IFNγ release assays (IGRA) for another 2 years (3). Individuals were classified as concordant negative resisters (RSTR) or concordant positive latent tuberculosis infection (LTBI). All participants were at least 15 years old at the time of retracing, HIV-negative, and gave written, informed consent, approved by the institutional review boards of their associated institution. Cryopreserved peripheral blood mononuclear cells (PBMC) and plasma from a subset of retraced individuals were used here.

### Cell culture and DNA extraction

Cryopreserved PBMCs were thawed, washed, and resuspended at 2E6 cells/mL in RPMI-10 supplemented with M-CSF (50 ng/mL), then rested overnight in 6-well non-TC treated dishes. CD14+/CD16+ monocytes were isolated by negative selection magnetic bead column purification (Pan Monocyte Isolation Kit, Miltenyi Biotec). For chromatin accessibility (ATAC-seq), 5x10E4 monocytes were removed and lysed in resuspension buffer (10 mM Tris-HCl pH 7.4, 10 mM NaCl, 3 mM MgCl_2_) plus 0.1% NP40, 0.1% Tween-20, and 0.01% digitonin. Cells were lysed on ice for 3 minutes and then washed in resuspension buffer plus 0.1% Tween-20. Nuclei were centrifuged for 10 min at 500xg and 4°C. Supernatant was removed and nuclei were resuspended in 50 μL of transposition mix (Tagment DNA Buffer, 0.1% Tween-20, 0.05% digitonin, 2.5 µL Tagment DNA Enzyme). Transposition reactions were incubated for 30 minutes at 37°C with shaking at 10E3 RPM. DNA fragments were purified with MinElute Reaction kit according to the manufacturer’s instructions (Qiagen). For methylation, monocytes were plated at 5x10E5 per well in RPMI-10 supplemented with M-CSF. After 24 hours, media was removed, and genomic DNA was isolated using Quick-gDNA MiniPrep kit according to the manufacturer’s instructions (Zymo Research).

### Chromatin accessibility

DNA from unstimulated monocytes was amplified by PCR for 5 cycles with indexing primer and barcoded primers. Amplicons were purified with the Agencourt AMPure XP Purification Kit (Beckman Coulter) and sequenced using 50 bp paired-end Tn5 transposase-accessible chromatin sequencing (ATAC-seq) on a NovaSeq6000. Sequence quality was assessed using FastQC (v0.11.8 (45)), and sequencing adapters were removed using AdapterRemoval (v2.3.1 (46)). Reads were aligned to the human genome (GRCh38) using STAR (v2.7.5 (47)), and alignments were assessed with Picard (v2.33.3 (48)) and samtools (v1.10 (49)). Peaks were called from alignments using Genrich (v0.6 (50)) with PCR duplicates removed and using the default ATAC-seq mode. Further data cleaning was completed in R (v4.0.2 (51)). Consensus peaks across samples were determined using ChIPQC (v1.24.1 (52)). Blacklisted (5421 (53)), mitochondrial (1), and rare peaks (32537 present in < 10% of samples) were removed. Nucleosome-free reads in peaks were quantified with Rsubread (v2.2.4 (54)) and filtered to peaks on autosomal chromosomes with > 0.1 counts per million (CPM) in at least 10% of samples. This resulted in 40919 peaks and 1.3 million ± 1.1 million s.d. reads per sample. Reads were TMM normalized with edgeR (v3.30.3 (55)) and log2 CPM normalized with limma (v3.44.3 (56)). Peaks were annotated to the human genome using ChIPseeker (v1.24.0 (57)) and the UCSC known gene track for GRCh38 (58).

### Methylation

DNA from unstimulated monocytes was bisulfate treated using the EZ-96 DNA Methylation kit (Zymo Research). Bisulfite conversion was confirmed by PCR using Universal Methylated Human DNA Standard with hMLH1 Primers (Zymo Research). Converted DNA was then applied to an Illumina Infinium MethylationEPIC850 BeadChip (865918 probes) and sequenced on a NovaSeq6000. Data cleaning and analysis were performed using ChAMP (v2.18.3 (59)) in R (51). Probes were filtered (59)to remove poor-quality (23907), non-CpG (2860), XY chromosome (16424), and SNP adjacent probes (94347). Beta values for the remaining 728380 probes were normalized using functional normalization, chip bias was removed using Combat, and values were log2 transformed to M values. Probes were annotated to the human genome (hg19) within ChAMP and ported to GRCh38 (60).

### Kinship

Kinship was determined as previously described (9). Briefly, genotypes were determined using the Illumina MEGA^EX^ array or Infinium OmniExpress BeadChip as previously reported (44). In PLINK2 (61), SNPs present in both arrays were filtered by Hardy-Weinberg Equilibrium (P < 1x10E-6), minor allele frequency (MAF > 0.05), call rate (> 0.95), and linkage disequilibrium (LD R^2^ < 0.1 in 50 bp windows with a 5 bp slide). Pairwise kinship was calculated using the robust King method for identity-by-descent (IBD, SNPRelate v1.22.0 (62)) and a genetic relationship matrix (GRM, GENESIS v2.18.0 (63)) calculated from these 63812 filtered SNPs.

### Differential analyses

Differentially methylated probes (DMP) and differentially methylated regions (DMR) were determined in 29 RSTR and 29 LTBI samples with complete methylation and kinship data. Differentially accessible regions (DAR) were determined for 29 RSTR and 31 LTBI with complete chromatin accessibility and kinship data. DMP and DAR were assessed with kimma (v1.4.4 (64)) using a linear mixed effects model of RSTR vs LTBI corrected for age, sex, and kinship. Genetic kinship was included to account for closely related household contacts, and it improved model fit as assessed by sigma (Figure S1A). Sex and age were included as co-variates as they improved model fit for some sites (Figure S1B) and were significant for many sites (Figure S1C,D). Additional co-variates including body mass index (BMI), tuberculosis exposure score (3), and BCG vaccination scar were removed due to missing data, lack of significance, or no improvement in model fit assessed by residual sigma.

DMP model estimates were then used to determine differentially methylation regions (DMR) in DMRcate (v2.2.1 (65)) with settings recommended in (66) (min probes = 2, lambda = 500 bp, scaling = 5). DMR were assessed at FDR < 1E-70, a slight relaxation of DMRcate’s default cutoff of 8.16E-85, which was determined based on the rate of DMP within the data set.

### Gene enrichment

Epigenetic sites were annotated to hg38 genes based on overlap with intron, exon, untranslated region (UTR), or promoters defined as within 1500 bp of the transcription start site (TSS1500). DAR were annotated using ChIPseeker, DMP using the Infinium hg38 manifest (60), and DMR using DMRcate. Genes were queried for enrichment in Broad MSigDB (23) Hallmark (H) and gene ontology biological process (C5 GO BP) gene-sets using hypergeometric enrichment with SEARchways (v1.0.0). Gene-sets with FDR < 0.2 and > 4% enrichment (k/K) were considered significant. Gene ontology term similarity was visualized by semantic similarity using rrvgo implementing GOSemSim (67). DAR genes were not assessed due to the small number of associated genes.

### Gene expression

Previously, Ugandan RSTR and LTBI monocyte gene expression was profiled with and without 6-hour Mtb infection (9). Differentially expressed genes (DEGs) were defined for the Mtb:RSTR interaction term in a model of Mtb*RSTR corrected for age, sex, sequencing batch, and genetic kinship. This original model was used for ABCA1 expression in the present study. Since several methylation genes of interest did not meet the original study’s gene expression cutoff (1 CPM in at least 5% of samples), an additional targeted analysis of these genes was performed. Here, all genes with > 0 CPM in at least 1 sample were obtained and log2 counts per million (CPM) were calculated. Genes of interest (APOC3, CIT, KCNQ1, PLA2G3, SHANK2) were modeled using the same interaction model as the original study, and significant genes were defined at Benjamini-Hochberg FDR < 0.2. In addition, targeted gene-set enrichment analysis (GSEA (68)) was performed for the 11 MSigDB gene-sets that were significantly enriched in methylation genes. Gene-level fold change estimates were extracted from the original linear mixed effects model for the four Mtb:RSTR contrasts (e.g. Mtb vs media in RSTR, RSTR vs LTBI in media, etc). Significant enrichment using GSEA was defined at P < 0.05.

### Monocyte-derived macrophages (MDM) stimulated with IFN

Additional analyses were performed with a previously published RNA-seq data set of human MDM stimulated with IFNα8, IFNβ, IFNε, or IFNγ (24). Briefly, MDMs isolated from 5 healthy donors were treated with recombinant interferons for 6 hours. Total RNA was extracted and sequenced as described (24). In this study, linear mixed effects model fold change estimates of IFN-stimulated expression vs unstimulated controls were used in GSEA (68) of the 11 MSigDB gene-sets that were significantly enriched in methylation genes. Significant enrichment using GSEA was defined at P < 0.05.

### Cholesterol efflux capacity

Cholesterol efflux capacity (CEC) was measured in serum from 41 RSTR and 43 LTBI. CEC of serum HDL was quantified using J774 murine macrophages (ATCC) to measure total CEC and baby hamster kidney fibroblast (BHK) cells transfected with a mifepristone inducible ABCA1 transporter to measure ABCA1-specific CEC as previously described (69, 70). In short, serum from RSTR and LTBI was reconstituted from freshly thawed plasma, and polyethylene glycol (20% PEG8000, 2:5 v/v) was added to precipitate apoB-containing (non-HDL) lipoproteins. After centrifugation, supernatants containing HDL (serum HDL) were incubated for 4 hours with unstimulated or stimulated cells labeled with radiolabeled [^3^H]-cholesterol (71). For J774 macrophages, the cells were stimulated with cAMP to activate the cells including induction of ABCA1 expression. For BHK cells, stimulation was completed with 0.01 μM mifepristone to specifically induce expression of the ABCA1 transporter. Thus, the J774 cell model captures overall efflux of cholesterol through multiple pathways, including ABCA1, while the ABCA1-BHK model specifically measures cholesterol efflux through the ABCA1 pathway.

For both J774 and BHK cells, HDL CEC (percent total cholesterol effluxed from cells to HDL) was determined as the ratio of radio-labeled cholesterol in the media at the end of the incubation relative to the total in the system (media + cell pellet). For BHK cells, ABCA1-specific efflux was calculated as the difference in the percent efflux between stimulated cells more highly expressing ABCA1 versus unstimulated cells. Total CEC and ABCA1-specific CEC were compared in RSTR and LTBI in a linear mixed effects model corrected for age, sex, and BMI using kimma (64). Significance was evaluated at P < 0.05.

### HDL particle concentration and size

HDL was assessed for the same participants as CEC. Total lipoprotein fraction was separated from plasma proteins by a single step density ultracentrifugation of EDTA plasma as described previously (72, 73). HDL particle concentration and size were quantified by calibrated differential ion mobility analysis on a differential mobility analyzer (DMA) (TSI Inc., MN) as described previously (72, 73). Six HDL subspecies (extra small, small, medium, medium large, large, extra-large) were fitted to the DMA profiles by unsupervised, iterative curve-fitting using Fityk (74). Because DMA directly quantifies the number of particles, areas under the curve fitted for each subspecies were directly converted to HDL particle concentration using a calibration curve constructed with a protein standard. For total HDL particle concentration, intra-day and inter-day coefficients of variation (CV) were < 10%. For individual species, CV were < 10% with the exception of xsHDL and sHDL (14.8 and 18.5%, respectively). HDL concentration and size were compared in RSTR and LTBI in a linear mixed effects model corrected for age, sex, and BMI using kimma (64). Significance was evaluated at FDR < 0.2.

### Data availability

Access to raw epigenetic and transcriptomic data for the Uganda cohort is available through the NCBI database of Genotypes and Phenotypes (dbGaP) Data Browser (https://www.ncbi.nlm.nih.gov/gap/) under accession 002445.v3.p1 but first must be approved by the data access committee (DAC) for the study site (see Supplemental Methods in (9)). Data related to the MDM-IFN experiment are available in the Gene Expression Omnibus (GEO) GSE236156. Scripts for this manuscript are available at https://github.com/hawn-lab/RSTR_epigenetics_public

## ACKNOWLEDGEMENTS

We thank the individual study participants of the Kawempe Community Health Study, study coordinators and the clinical and research staff including LaShaunda Malone, Keith Chervenak, Marla Manning, Dr. Alphonse Okwera, Dr. Moses Joloba, Hussein Kisingo, Sophie Nalukwago, Dorcas Lamunu, Deborah Nsamba, Annet Kawuma, Saidah Menya, Joan Nassuna, Joy Beseke, Michael Odie, Henry Kawoya, and Bonnie Thiel.

