## Supplemental figures for "Epigenetic programming of host lipid metabolism associates with resistance to TST/IGRA conversion after exposure to *Mycobacterium tuberculosis*"

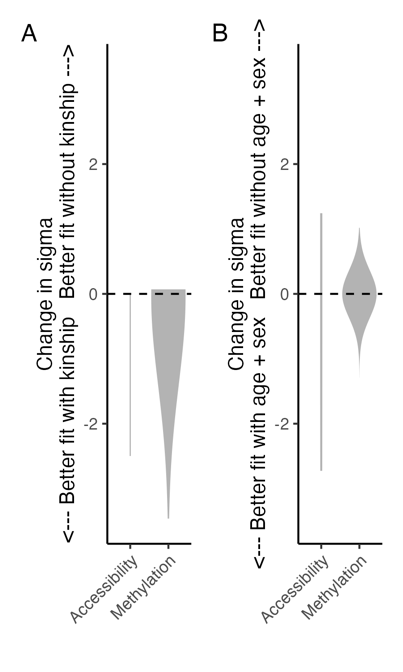

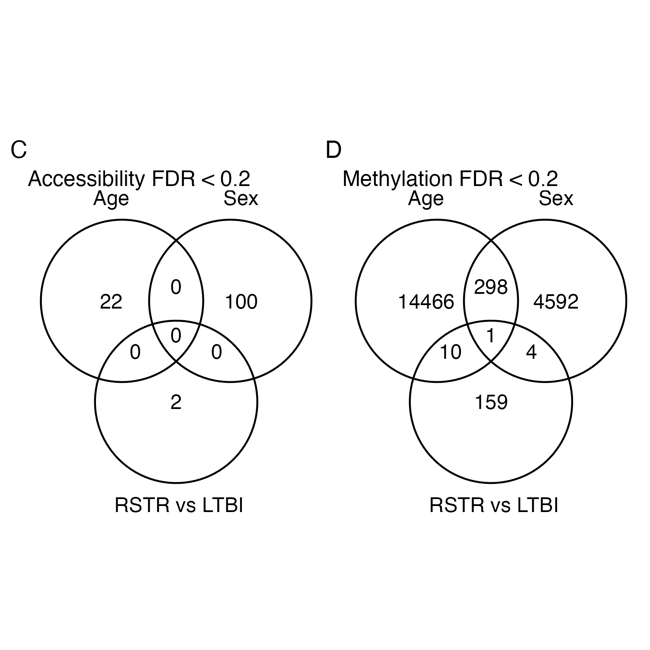

**Figure S1. Model fitting for methylation and chromatin accessibility data.** Accessibility peaks and methylation probes were assessed for differences between RSTR and LTBI corrected for kinship, age, and/or sex. Model sigma was compared between models (A) with and without kinship as well as (B) with and without age + sex for each peak/probe. Plots are violin distributions across all probes/peaks. Kinship improved model fit with smaller sigma for the majority of sites in accessibility (62%) and methylation (78%) data sets. Age and sex improved model fit for some sites in accessibility (39%) and methylation (51%) data. In addition, age and sex were significant for some sites for both (C) accessibility and (D) methylation.

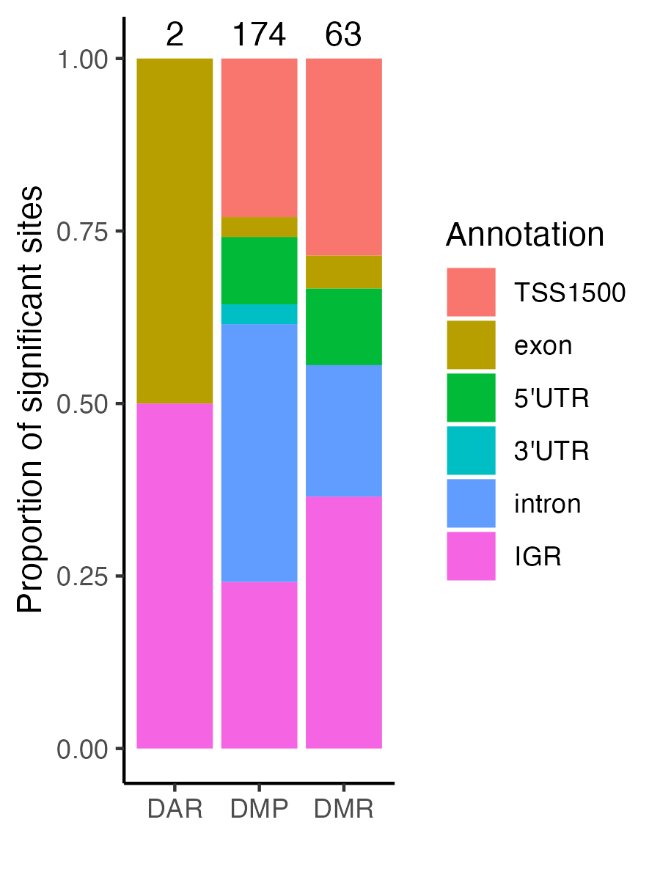

**Figure S2.** **Annotation of significant epigenetic regions.** Significant differentially accessible regions (DAR), differentially methylated probes (DMP), and differentially methylated regions (DMR) were annotated to genomic location features. In total, 2 DAR, 174 DMP, and 63 DMR were annotated to genes. Annotations included within 1500 bp of the transcription start site (TSS1500), within a gene (exon, intron), within an untranslated region (5’UTR, 3’UTR), and non-gene-specific intergenic regions (IGR). DMRs with multiple annotations were assigned to their topmost annotation as defined by the legend order (TSS1500 > exon > 5’UTR > 3’UTR > intron > IGR).

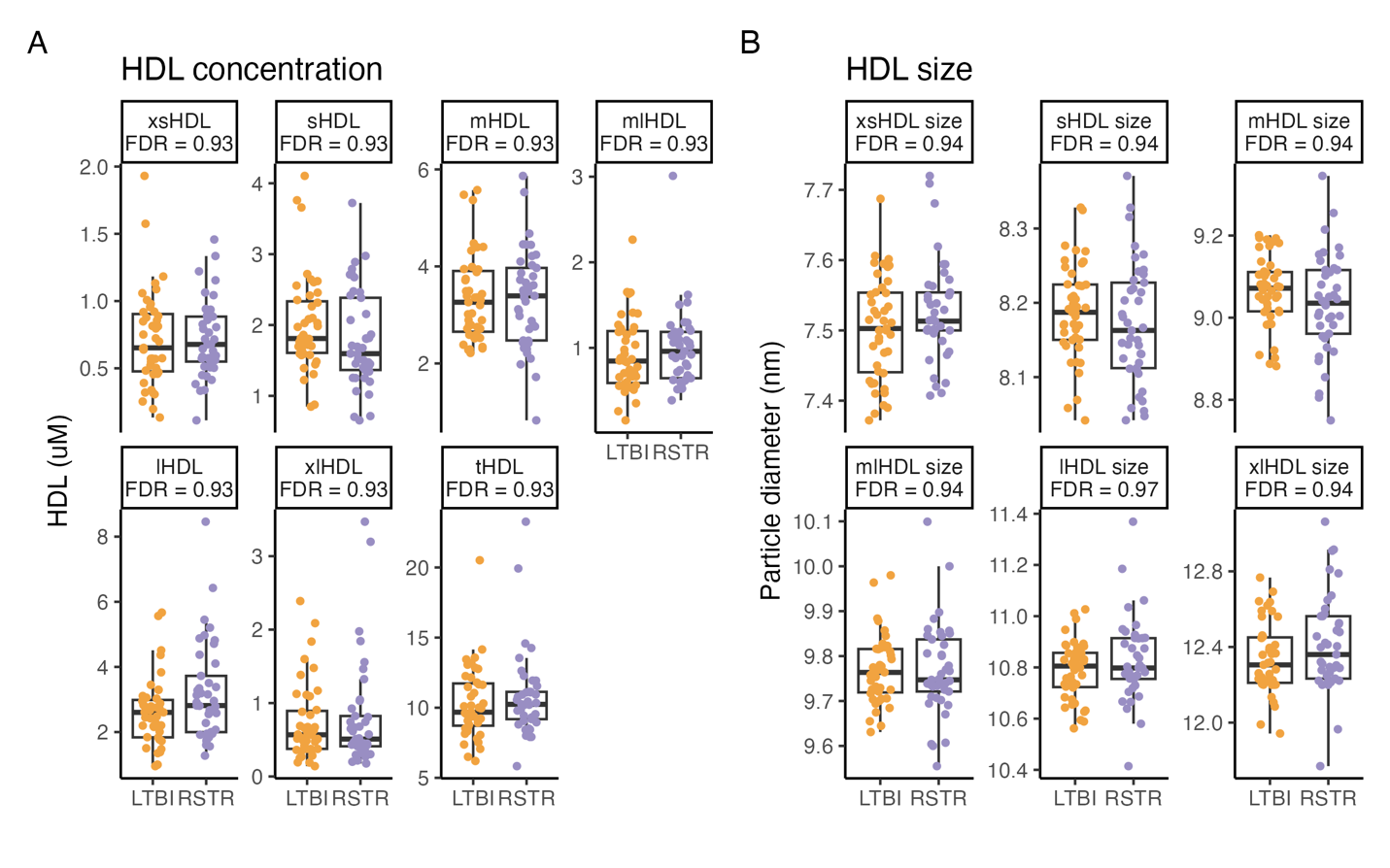

**Figure S3.** **HDL quantification, size, and efflux.** HDL was isolated from plasma from RSTR and LTBI subjects and analyzed further for (A) HDL concentration (uM) and (B) HDL size (nm) separated by major size classes (xs to xl). xs: extra-small, s: small, m: medium, ml: medium-large, l: large, xl: extra-large, t: total

**SUPPLEMENTAL TABLES**

**Table S1. Differentially accessible regions, differentially methylated probes, and differentially methylated regions in RSTR vs LTBI monocytes.** Differentially accessible regions (DAR, FDR < 0.2), differentially methylated probes (DMP, FDR < 0.2), differentially methylated regions (DMR, FDR < 1E-70), and probes in DMRs (DMR_probe) from RSTR vs LTBI comparison with genomic region annotations. Genomic positions and gene annotations were determined using hg38. Annotations were simplified to groups for in genes (intron, exon), within 1500 of the transcription start site (TSS1500), in untranslated regions (3’UTR, 5’UTR), or intergenic region / no annotation (IGR). CHR: chromosome; start, end: genomic position in bp; LTBI_mean, RSTR_mean: Mean normalized log2 CPM (DAR) or mean normalized M values (DMP, DMR) for each group

**Table S2. Hypergeometric mean pathway enrichment analysis of differentially methylated probes and regions in RSTR vs LTBI monocytes.** Hypergeometric mean pathway enrichment analysis was performed with genes annotated to 174 DMP or 63 DMR. Gene sets were tested from MSigDB Hallmark and gene ontology (C5 GO). Only DMR genes in GO resulted in significant gene sets (FDR < 0.2, k/K > 4%). GOID: gene ontology identifier; group_in_pathway: number of significant genes in gene set (k); size_pathway: number of genes in gene set (K).

**Table S3. DMR annotation.** Differentially methylated region (DMR) annotations including genes, total probes, probe-level RSTR vs LTBI fold change, and mean region-level RSTR vs LTBI fold change. Fold changes are as in Figure 2B.

**Table S4.** **Gene set enrichment analysis (GSEA) of RNAseq.** Gene set enrichment analysis (GSEA) was performed for pathways that were significantly enriched for DMR annotated genes. Fold change estimates were used from mixed effects models run on one of two RNAseq data sets. (MDM-IFN-GSEA) GSEA for IFN stimulation. MDMs from healthy donors were stimulated with type I and II interferons for 6 hrs. Gene expression was modeled for IFN vs media. (MTB-RSTR-GSEA) GSEA for Mtb and RSTR contrasts. Monocytes from RSTR and LTBI were infected with Mtb for 6 hours, and gene expression was modeled for the interaction of Mtb and RSTR status (Mtb:RSTR) corrected for age, sex, kinship, and sequencing batch. Contrast fold changes were compared for Mtb infection vs media within RSTR or LTBI as well as RSTR vs LTBI within media or Mtb infected groups.

**Table S5.** **HDL linear models.** HDL was isolated from plasma from RSTR and LTBI subjects. HDL quantification (HDL), mean HDL size (HDL_size), and cholesterol efflux capacity (CEC) in J774 and BHK cells (HDL_efflux) were modeled for RSTR vs LTBI corrected for age, sex, and body mass index (BMI). CEC was measured with HDL incubated for 4 hours with [^3^H]-cholesterol loaded cells that were or were not stimulated to increase CEC. CEC was calculated as percent cholesterol in media versus total (media + cell pellet). J774 macrophages were stimulated with cAMP to increase CEC. BHK cells were stimulated with mifepristone to induce ABCA1 expression. xs: extra-small, s: small, m: medium, ml: medium-large, l: large, xl: extra-large, t: total
